# Cell type- and stage-specific expression of Otx2 is coordinated by a cohort of transcription factors and multiple *cis*-regulatory modules in the retina

**DOI:** 10.1101/2019.12.19.882969

**Authors:** Candace Chan, Nicolas Lonfat, Rong Zhao, Alexander Davis, Liang Li, Man-Ru Wu, Cheng-Hui Lin, Zhe Ji, Constance L. Cepko, Sui Wang

## Abstract

Transcription factors (TFs) are often used repeatedly during development and homeostasis to control distinct processes in the same and/or different cellular contexts. Considering the limited number of TFs in the genome and the tremendous number of events that need to be regulated, re-use of TFs is an advantageous strategy. However, the mechanisms that control the activation of TFs in different cell types and at different stages of development remain unclear. The neural retina serves as a model of the development of a complex tissue. We used this system to analyze how expression of the homeobox TF, Orthodenticle homeobox 2 (Otx2), is regulated in a cell type- and stage-specific manner during retinogenesis. We identified seven *Otx2 cis*-regulatory modules (CRMs), among which the O5, O7 and O9 CRMs mark three distinct cellular contexts of Otx2 expression. These include mature bipolar interneurons, photoreceptors, and retinal progenitor/precursor cells. We discovered that Otx2, Crx and Sox2, which are well-known TFs regulating retinal development, bind to and activate the O5, O7 or O9 CRMs respectively. The chromatin status of these three CRMs was found to be distinct in vivo in different retinal cell types and at different stages, as revealed by ATAC-seq and DNase-seq analyses. We conclude that retinal cells utilize a cohort of TFs with different expression patterns, and multiple CRMs with different chromatin configurations, to precisely regulate the expression of Otx2 in a cell type- and stage-specific manner in the retina.

## Introduction

In the mammalian genome, there are approximately 1500 transcription factors (TFs) (Zhou et al. 2017; Vaquerizas et al. 2009), which bind to specific DNA sequences, within *cis*-regulatory modules (CRMs), to control gene expression and regulate almost every aspect of life. Many are used repeatedly, and in a variety of combinations, to regulate distinct developmental and homeostatic events. For instance, a handful of TFs, including Pax6, Ptf1a, Sox9, Hnf6 (Onecut1) and NeuroD1, regulate pancreatic development as well as retinal development (Bastidas-Ponce et al. 2017; Emerson et al. 2013; Poché et al. 2008, 9; Ohsawa and Kageyama 2008). Moreover, within the same organ, TFs can be expressed in mitotic cells, newly postmitotic cells, and mature cells to regulate development as well as function. The reuse of a limited pool of TFs in different cellular contexts likely evolved to maximize their utility in driving the evolution of an amazingly diverse set of cell types and functions. A full definition of how specificity and function are created through the redeployment of TFs is an area of interest in many systems.

The murine neural retina has served as a model system for studying the development of a complex mammalian tissue (Cepko et al. 1996). Access to the developing retina using in vivo electroporation has provided a strong platform for studies of gene regulation, as well as cell fate determination (Matsuda and Cepko 2007, 2004). The developmental role and regulation of Otx2, a homeobox TF that is important in multiple aspects of retinal development, has been the subject of several such studies. Otx2 is expressed in retinal progenitor cells (RPCs), the mitotic cells of the retina. RPCs that express Otx2 are a subset of those that are about to produce postmitotic daughter cells (Muranishi et al. 2011; Trimarchi et al. 2008). In addition, Otx2 is expressed in mature photoreceptor cells, the rods and cones, as well as in mature bipolar interneurons (Koike et al. 2007; Fossat et al. 2007). Otx2 has been shown to be required for the genesis of photoreceptor cells, bipolar cells, as well as a type of interneuron, horizontal cells, which do not express Otx2 (Koike et al. 2007; Nishida et al. 2003; Sato et al. 2007). The dosage of Otx2 is important for the determination of rods vs. bipolar interneurons (Wang et al. 2014), as well as for survival and activity of mature photoreceptors, bipolar cells, and horizontal cells (Housset et al. 2013; Béby et al. 2010; Bernard et al. 2014). Over-expression of *Otx2* in neonatal mouse retinas in vivo, when rods and bipolar cells are being generated, was found to increase the production of bipolar cells at the expense of rod photoreceptors. Knocking down *Otx2* levels via shRNA led to the formation of more rods and fewer bipolar cells (Wang et al. 2014). These and other studies (Martinez-Morales et al. 2001) establish *Otx2* as a key gene in the development of several ocular tissues and retinal cell types, posing interesting questions about how it is regulated to achieve this variety of outcomes.

Several studies of the regulation of *Otx2* during retinal development have been carried out. One CRM, “EELPOT”, can recapitulate Otx2 expression in embryonic mouse retinas (Muranishi et al. 2011). It is positively regulated by Rax and negatively regulated by the Notch signaling pathway. Another CRM, designated ECR2, recapitulates Otx2 expression in a subset of neonatal RPCs and their newly postmitotic daughters (Emerson and Cepko 2011). ECR2 has not been characterized regarding its TF binding sites (TFBSs) or cognate TFs. In addition, three DNaseI hypersensitive genomic fragments (DHS-2, DHS-4 and DHS-15) flanking the *Otx2* gene have been shown to be active in the neonatal mouse retina (Wilken et al. 2015), though their upstream TFs and expression patterns were not characterized. Despite these previous studies, the CRMs that regulate Otx2 expression in mature photoreceptor and bipolar cells are currently unknown. In addition to regulation in these various cell types, the regulation of expression levels of Otx2 is of interest. As rod and cone photoreceptors differentiate, the Otx2 level decreases, while it increases in bipolar cells, and disappears in differentiating horizontal cells (Trimarchi et al. 2008; Koike et al. 2007; Fossat et al. 2007; Baas et al. 2000). Dose is also important for the survival and function of mature retinal cells, as alleles that reduce the level and activity of Otx2 lead to retinal degeneration (Bernard et al. 2014).

To elucidate the regulation of *Otx2* in different cellular contexts, we systematically searched for candidate CRMs for *Otx2* using the DNaseI hypersensitivity of genomic regions within 350 kb of the *Otx2* gene. Candidates were tested for CRM activity, and seven novel *Otx2* CRMs were identified. Three of these CRMs, designated O5, O7 and O9, were found to drive strong expression in the postnatal retina in vivo. O5 drives expression primarily in mature bipolar cells. O7 drives expression in mature rods, while O9 drives expression in developing neonatal cells. These three CRMs thus recapitulate Otx2 expression in distinct cell types and at different stages. We further explored the TF binding sites (TFBSs) and TFs that regulate these CRMs, and discovered that Otx2, Crx and Sox2 can bind to and activate O5, O7, or O9 respectively. We also examined the endogenous chromatin status of O5, O7 and O9 CRMs in different retinal cell types, and found that these three CRMs show different chromatin states in different cell types. We conclude that the expression of Otx2 in different retinal cellular contexts is not regulated by common regulators. The chromatin accessibility and presence of particular TFs together ensure the expression of Otx2 in a cell type- and stage-specific manner in the retina.

## Results

### Identification of active CRMs for the *Otx2* gene based on chromatin status

CRMs have been nominated using a variety of methods and features. Chromatin accessibility and histone modifications are two such features (Klemm et al. 2019). The UW ENCODE project created high quality, genome-wide mapping of DNaseI hypersensitive sites (DNaseI HS) for major tissues and cell types in mice and humans, including the mouse neural retina (The ENCODE Project Consortium 2011; Vierstra et al. 2014). We systematically evaluated 25 DNaseI HS fragments, spanning an approximately 350Kb region surrounding the *Otx2* locus, for CRM activity in the retina. Each DNaseI HS fragment was amplified and cloned into the Stagia3 reporter vector, which expresses GFP and Alkaline Phosphatase (AP) when an active CRM is inserted upstream of a minimal TATA promoter (Billings et al. 2010). When tested in newly explanted retinas ex vivo, seven of the DNaseI HS regions (O5, O7, O9, O10, O11, O15, and O20) drove significant expression of the reporter gene (Figure 1). Although the precise sequences of the three CRMs reported by Wilken et al. are not available, based on the approximate coordinates, the O5 and O9 CRMs may overlap with their DHS-2 and DHS-4.

**Figure 1.**
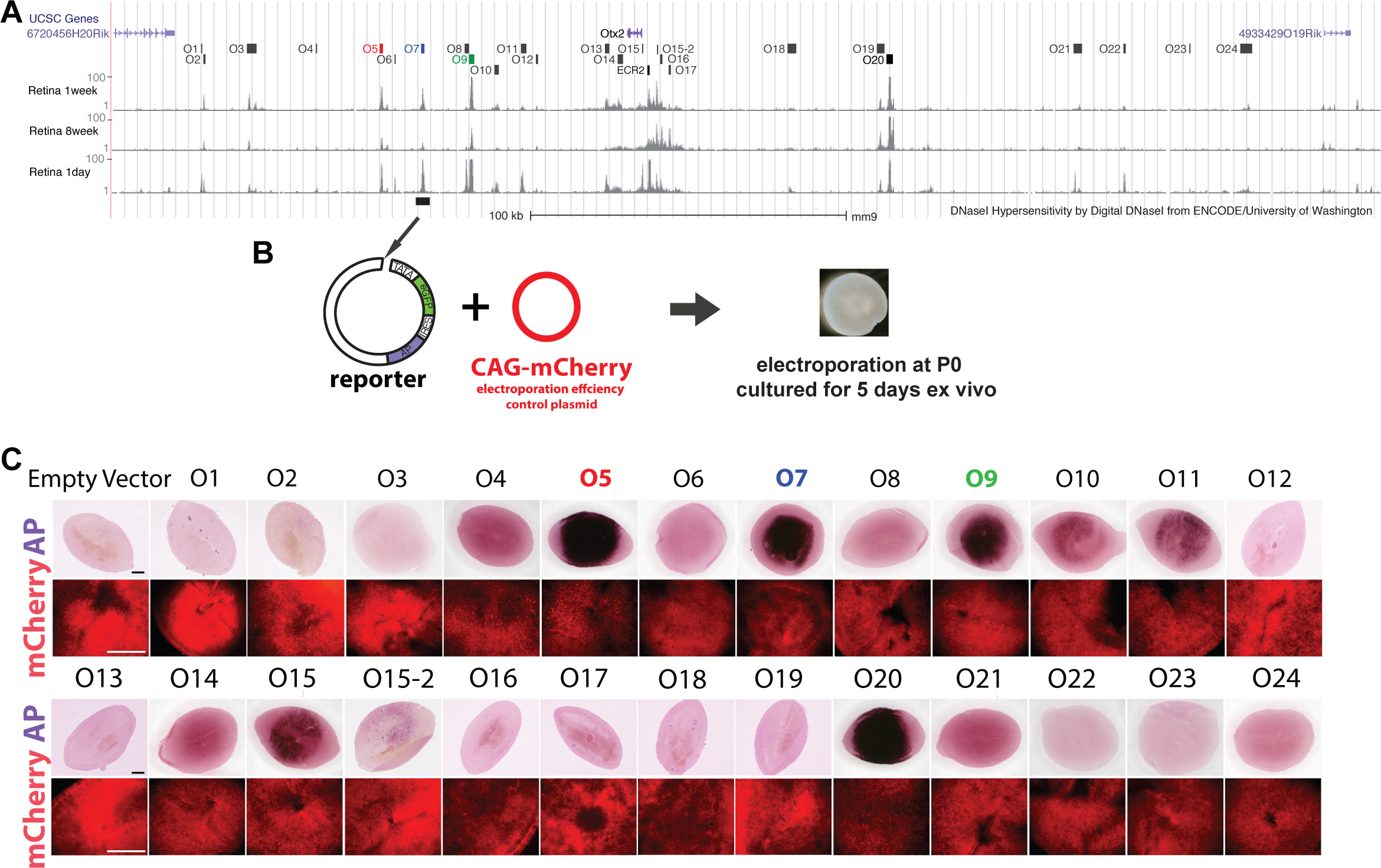
Assay of DNaseI hypersensitive regions near *Otx2* gene for CRM activity in the retina. **(A)** Twenty-five DNaseI hypersensitive regions (The ENCODE Project Consortium 2011; Vierstra et al. 2014), O1-O24, flanking the *Otx2* gene are highlighted. **(B)** Individual DNaseI hypersensitive regions were cloned into the Stagia3 reporter plasmid, and electroporated into P0 mouse retinas. The retinas were cultured as organ cultures for 5 days ex vivo. The CAG-mCherry plasmid served as the electroporation efficiency control. **(C)** Seven out of the 25 DNaseI hypersensitive regions drove AP (Alkaline phosphatase) expression. The O5, O7 and O9 CRMs (highlighted), became the focus of this study. Scale bar: 0.5mm.

We tested the activity of these seven CRMs in vivo by electroporating individual reporter plasmids into mouse retinas at postnatal day 0 (P0) (Figure 2). Electroporation tends to result in expression in RPCs and their postmitotic progeny, which in the neonatal period become primarily rods and bipolar cells, with a small number of amacrine interneurons and Müller glial cells (Matsuda and Cepko 2007; Young 1985). Plasmids can be retained in these mature cell types, allowing for a read out of activity in multiple cell types and their RPCs. Retinas were thus harvested at P3 or P14 to examine the activities of these CRMs in developing and mature retinal cells, respectively. The O10, O11, O15 and O20 CRMs were found to drive GFP expression in vivo, but were relatively weak (data not shown), and were not investigated further.

**Figure 2.**
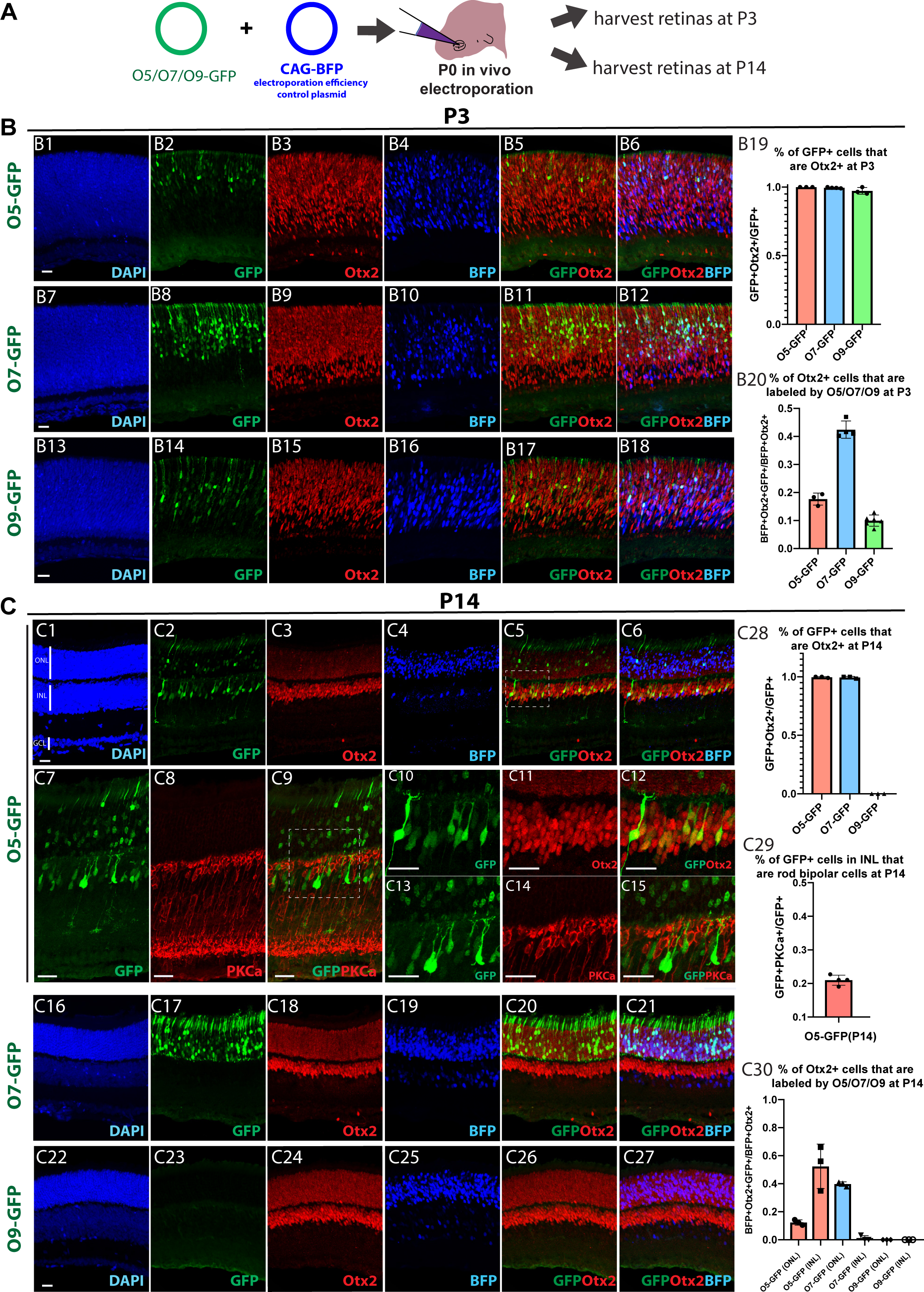
O5, O7 and O9 CRMs drove cell type- and stage-specific expression in the retina in vivo at postnatal stages. **(A)** The O5, O7 or O9 reporter plasmids (O5-GFP, O7-GFP or O9-GFP) were electroporated into P0 mouse retinas in vivo, together with the CAG-BFP plasmid, which served as the electroporation efficiency control. The retinas were harvested at P3 or P14. **(B)** The activity of the O5 (B1-B6), O7 (B7-B12) and O9 (B13-B18) CRMs in the retina in vivo at P3. The retinal sections were stained with anti-Otx2 antibody and DAPI. The percentage of GFP+ cells that were Otx2+ and the percentage of Otx2+ cells that were labeled by individual CRMs are quantified in B19 and B20 respectively. **(C)** The activity of the O5 (C1-C15), O7 (C16-C21) and O9 (C22-C27) CRMs in the retina in vivo at P14. The retinal sections were stained with anti-Otx2 (C1-C6, C10-C12, and C16-C27) or anti-PKC-a antibody (C7-C9 and C13-C15), and DAPI. C10-C12: High magnification views of the highlighted region in C5. C13-C15: High magnification views of the highlighted region in C9. The percentage of GFP+ cells that were Otx2+ at P14 was quantified (C28) for individual CRMs. The percentage of O5-GFP+ cells that were rod bipolar neurons was quantified (C29). The percentage of Otx2+ cells that were labeled by individual CRMs at P14 is shown in C30. Scale bar: 20um.

At P3, the O5, O7 and O9 CRMs were found to direct strong GFP expression, mimicking the endogenous expression of Otx2 (Figure 2B). More than 97% of the GFP+ cells were positive for Otx2 protein expression, as detected using immunohistochemistry (IHC) (Figure 2, B19). O5, O7 and O9 CRMs were active in approximately 16%, 42% and 10% of the electroporated cells that were Otx2+ (BFP+Otx2+ cells) at P3, respectively (Figure 2, B20). Co-electroporation of multiple CRMs driving different fluorescent proteins revealed that there was a significant number of cells simultaneously expressing the reporters driven by two or all three of these CRMs (Supplementary Figure 1). We estimated that O5, O7 and/or O9 CRMs were collectively expressed in approximately 50% of the Otx2+ cells at P3 (Supplementary Figure 1J).

As Otx2 has been shown to be expressed in the G2-M phase of the last cell cycle in a subset of RPCs (Trimarchi et al. 2008), the expression driven by the CRMs was examined in mitotic cells. First, to confirm that a subset of RPCs expresses Otx2, we examined the expression of Otx2 protein in cells labelled by EdU, a S-phase marker (Buck et al. 2008), or Ki67, a pan-cell cycle marker (Scholzen and Gerdes 2000). A small subset of Otx2+ cells were found to be EdU+ (<2%, 30min EdU pulse, Supplementary Figure 2A-C & G) or Ki67+ (< 15%, Supplementary Figure 2D-F & H). The O5 and O7 CRMs were found to be less active in EdU+ cells, while the O9 CRM preferentially marked the Otx2+ cells that were cycling at P3 (Supplementary Figure 2). These data indicate that the O5, O7 or O9 CRMs mark overlapping, but distinct, populations of cells that express Otx2 in the neonatal mouse retina, with O9 being the most active in mitotic cells.

At P14, when the retinal is nearly fully developed, Otx2 is expressed at higher levels in bipolar cells, and relatively lower levels in photoreceptor cells (Figure 2, C3). The O5 CRM was observed to mimic this pattern (Figure 2, C1-6). In the inner nuclear layer (INL), the majority of O5-GFP+ cells were found to be cone bipolar cells, as recognized by the characteristic axonal projection pattern (Behrens et al.). In addition, approximately 20% of O5-GFP+ cells in the INL were positive for PKCa (Figure 2, C7-15 & C29), a well-established marker of rod bipolar cells (Negishi et al. 1988; Shekhar et al. 2016). Given that the estimated cone bipolar to rod bipolar ratio is about 2.6:1 in mouse retina (Strettoi et al. 2010), the O5 CRM is biased towards labeling cone bipolar cells. In contrast, the activity of the O7 CRM was restricted to rod photoreceptors at P14 (Figure 2, C16-21), and the O9 CRM was negative at P14 (Figure 2, C22-30).

Taken together, the O5, O7 and O9 CRMs direct expression recapitulating the endogenous Otx2 expression in different retinal cell populations and at different developmental stages. The O9 CRM preferentially labels Otx2+ RPCs and their newly postmitotic daughters, and its activity is silenced in differentiated cell types. In mature retinas (P14 or later), the O5 CRM drives expression predominantly in bipolar cells and in a relatively small subset of rods, and the O7 CRM directs expression only in rod photoreceptors.

### The O5 CRM is required for expression of Otx2 in bipolar cells

In order to determine if the *Otx2* CRMs identified above are necessary for endogenous *Otx2* expression, we deleted them from the genome in vivo. We electroporated a Cas9 plasmid and appropriate single guide RNAs (sgRNAs) into the mouse retina in vivo for this purpose. Plasmids encoding sgRNAs targeting the 5’ or 3’ region of the O5 CRM, as well as O5-Cas9 (O5 CRM drove Cas9 expression) and CAG-mCherry plasmids, were co-electroporated into mouse retinas at P0 in vivo (Figure 3A-B). Previous studies have shown that there is a high efficiency of co-electroporating multiple plasmids into the same retinal cells in vivo (Matsuda and Cepko 2004, 2007). The retinas were harvested at P14 and the levels of endogenous *Otx2* transcripts were detected and quantified by single molecule fluorescent RNA in situ hybridization (smFISH) (Raj et al. 2008; Wang et al. 2012). The number of smFISH puncta had been found to correlate well with mRNA levels. We dissociated the retinas into single cell preparations to increase the accuracy of quantification. The electroporated cells that received the plasmid mix were labeled by mCherry (Figure 3C-F). Bipolar cells were labeled by IHC for Chx10 (Vsx2). The number of *Otx2* smFISH puncta in mCherry+Chx10+ bipolar cells with or without the O5 CRM deletion was quantified (Figure 3G). The number of bipolar cells with low *Otx2* mRNA levels was significantly increased when the O5 CRM was deleted, demonstrating that the O5 CRM is necessary for the wild type level of *Otx2* mRNA in bipolar cells.

**Figure 3:**
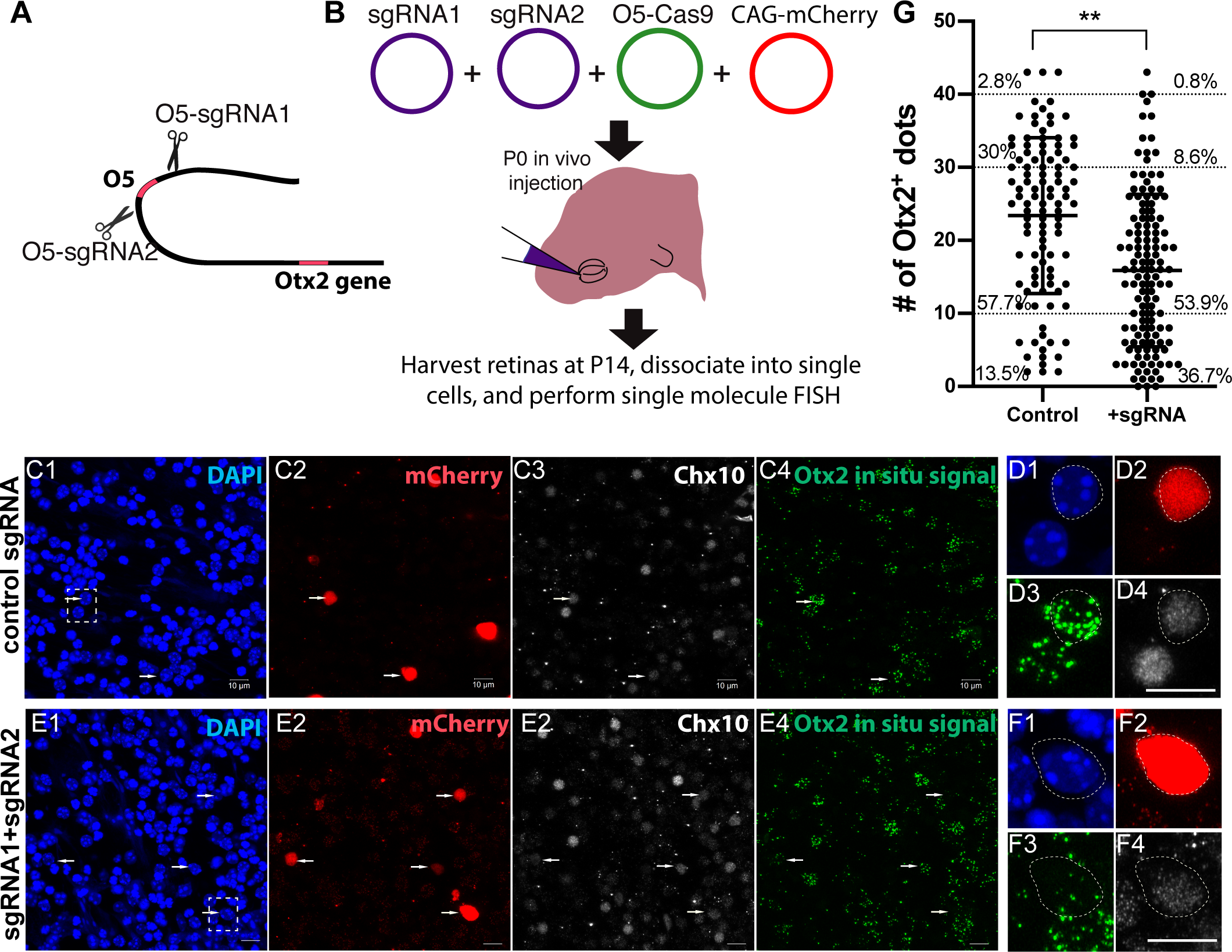
Deletion of the O5 CRM reduced *Otx2* transcript levels in bipolar cells. **(A&B)** The O5-sgRNA1 plasmid targets the 5’ end of the O5 CRM in the mouse genome, and the O5-sgRNA2 plasmid targets the 3’ of the O5 CRM. The O5-Cas9 plasmid expressed Cas9 under the control of the O5 CRM. The CAG-mCherry plasmid served as the electroporation efficiency control. These plasmids were co-electroporated into P0 retinas in vivo. At P14, retinas were harvested and dissociated into single cells. The transcript levels of *Otx2* in mCherry+Chx10 (Vsx2)+ bipolar cells were detected and quantified by smFISH. **(C & D)** The level of the Otx2 transcript in control retinal cells. mCherry marked the electroporated cells. Chx10 IHC labeled bipolar cells. The Otx2 transcript was detected by smFISH (Green signal). Each green dot in C4 represents one mRNA molecule. D1-D4 are high magnification views of the highlighted region in C1. White arrows: mCherry+Chx10+ bipolar cells. **(E & F)** The level of Otx2 transcripts in retinal cells that had the CRISPR constructs to delete the O5 CRM. F1-F4 are high magnification views of the highlighted region in E1. **(G)** Quantification of the Otx2 transcript levels. The Y axis represents the number of Otx2 transcripts (green dots) in individual bipolar cells based on smFISH (C4, D3, E4 and F3). Solid circles in the plot represents cells with each solid circle representing one cell. The percentage of cells that expressed different levels of Otx2 are indicated. **: P<0.01. Scale bar: 10um.

Assay for the endogenous role of the O9 CRM was not successful. Due to the extended time needed to accumulate significant levels of Cas9 protein, and the fact that the O9 CRM is only transiently active in Otx2+ progenitor/precursor cells, we were not able to delete the O9 CRM quickly enough to assay its endogenous activity. Deletion of the O7 CRM by CRISPR did not result in significant down regulation of *Otx2* mRNA levels (data not shown). Its function may be compensated by other *Otx2* CRMs, such as O5, which has activity in rods (Figure 2, C2).

### Identification of TFBSs and TFs for O5, O7 or O9 activity

To understand how O5, O7 or O9 CRMs are regulated, we searched for TFBSs and TFs that are required for activation of these CRMs. We first determined the minimal sequences within each CRM that are required for CRM activity. The O5, O7 or O9 CRMs were truncated into smaller fragments based on the positions of bioinformatically predicted TFBSs (MotifDb, Shannon 2015, Grant et al. 2011). Each fragment was individually cloned into the Stagia3 reporter and tested in mouse retinas ex vivo (Figure 4A, C & E). The O5-8, O7-1 and O9-12 elements were found to drive strong expression of the AP reporter gene. We then mutated individual predicted TFBSs on O5-8, O7-1 or O9-12 to further investigate the identity of their cognate TFs. Mutating the Otx2 binding site (Otx2 BS) completely abolished the activity of the O5-8 CRM (Figure 4B). Mutation of the Crx binding site (Crx BS) reduced the activity of O7-1 element (Figure 4D). Mutating the Sox2 binding site (Sox2 BS) reduced the activity of the O9-12 element (Figure 4F). The ability of Otx2, Crx or Sox2 to activate O5-8, O7-1 or O9-12 was further investigated in HEK293T cells in vitro, which do not normally activate the Otx2 CRMs. Otx2 was found to be sufficient to activate the wild type O5-8, but not the element with the Otx2 binding site mutation (Figure 4G). Crx and Sox2 behaved similarly on their respective elements (Figure 4H-I). To examine whether Otx2, Crx or Sox2 directly binds to O5, O7 or O9 CRMs, we performed electrophoretic mobility shift assays (EMSA) (Supplementary Figure 3). The biotin-labeled CRMs were used as DNA probes. Strong electrophoretic mobility bands were detected when the CRM probes were mixed with TF-enriched nuclear extracts. When un-labeled cold probe or antibody was included, the primary EMSA band disappeared, suggesting that the binding was specific. These results demonstrate that Otx2, Crx or Sox2 can regulate O5, O7 or O9 CRMs via direct binding, and likely contribute to the cell type- and stage-specific activation of these CRMs.

**Figure 4.**
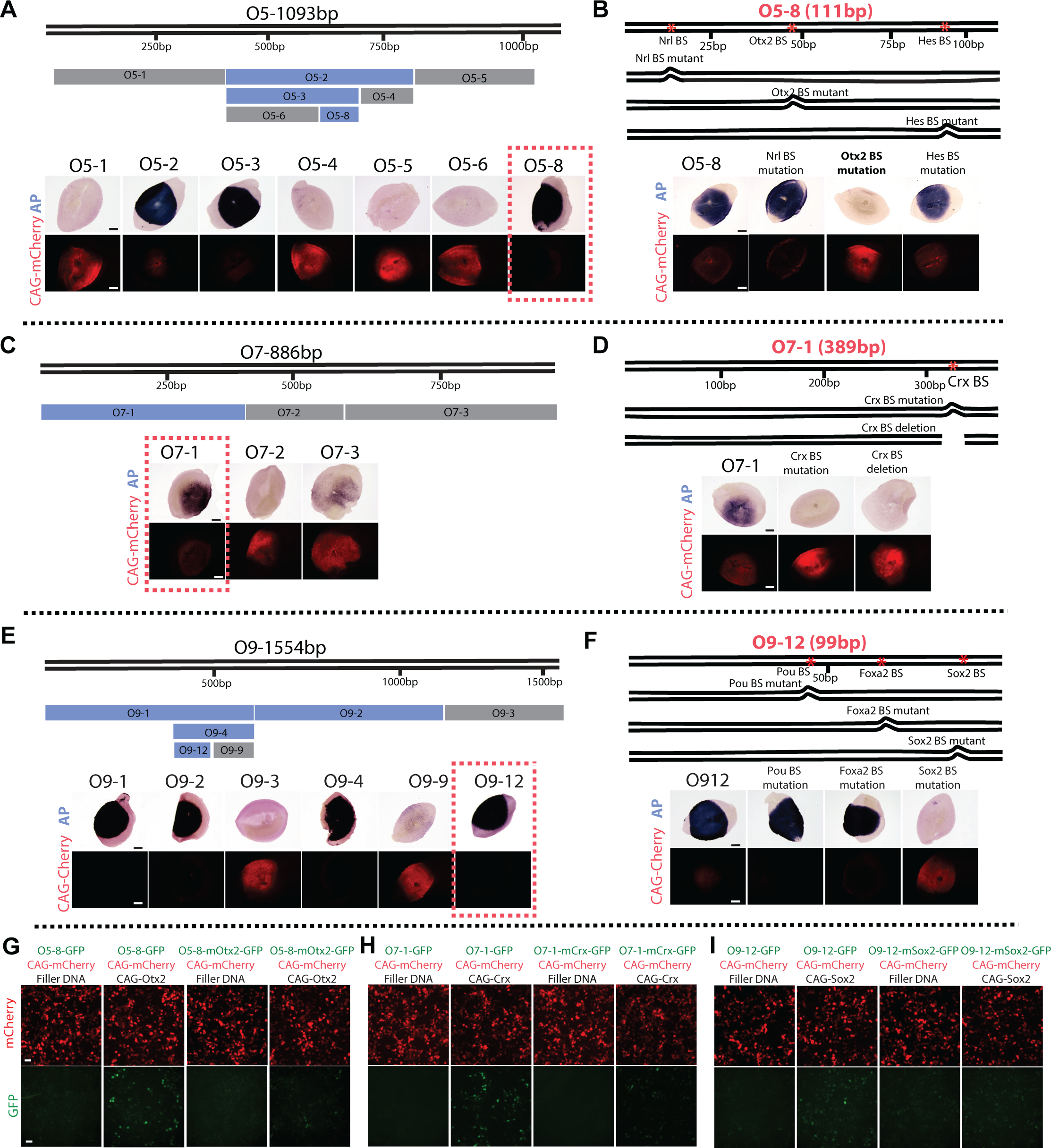
Otx2, Crx or Sox2 activated O5, O7 or O9 CRM respectively. **(A, C & E)** The minimal elements within O5 (A), O7 (C) or O9 (E), which were sufficient to drive reporter activity (AP signals), were discovered by creating deletion constructs of individual CRMs. Each CRM was truncated into smaller elements based on the availability of predicted TF binding sites. Each truncated element was incorporated into the Stagia3 reporter plasmid, and tested for its ability to drive AP expression in retinas ex vivo. CAG-mCherry plasmid was used as the electroporation efficiency control. Strong AP signals often absorbed the fluorescent mCherry signals (e.g. O5-2). Scale bar: 0.5mm. **(B, D & F)** The Otx2, Crx or Sox2 binding sites were required for the activity of O5-8 (B), O7-1 (D) or O9-12 (F) elements, respectively. The mutation or deletion of these binding sites abolished reporter activity. Scale bar: 0.5 mm. **(G, H & I)** Otx2, Crx or Sox2 were sufficient for reporter activity in HEK293T cells. The CAG-mCherry plasmid was the transfection efficiency control. In the O5-8-mOtx2-GFP plasmid, the Otx2 binding site was mutated. In the O7-1-mCrx-GFP plasmid, the Crx binding site was mutated. In the O9-12-mSox2-GFP plasmid, the Sox2 binding site was mutated. Scale bar: 50um.

To investigate if the expression patterns of Otx2, Crx and Sox2 match what would be predicted for their roles in regulating the *Otx2* CRMs, we checked their patterns of expression by IHC. At neonatal stages, O5-GFP+, O7-GFP+, or O9-GFP+ cells were positive for Otx2, Crx or Sox2 respectively (Figure 2B and Supplementary Figure 4). However, not all Otx2+, Crx+, or Sox2+ electroporated cells turned on O5, O7 or O9 CRM by P3. This could be due to the levels of the TFs, which might not be high enough in certain retinal cells. To test this possibility, we over-expressed Otx2, Crx or Sox2 in P0 retinas electroporated by the CRM reporter plasmids. This led to increased numbers of cells that turned on O5, O7 or O9 CRM plasmids within 24 hours (Supplementary figure 5). These data suggest that Otx2, Crx or Sox2 can activate O5, O7 or O9 CRM reporters at neonatal stages, and the levels of these TFs may be important for CRM activity.

**Figure 5.**
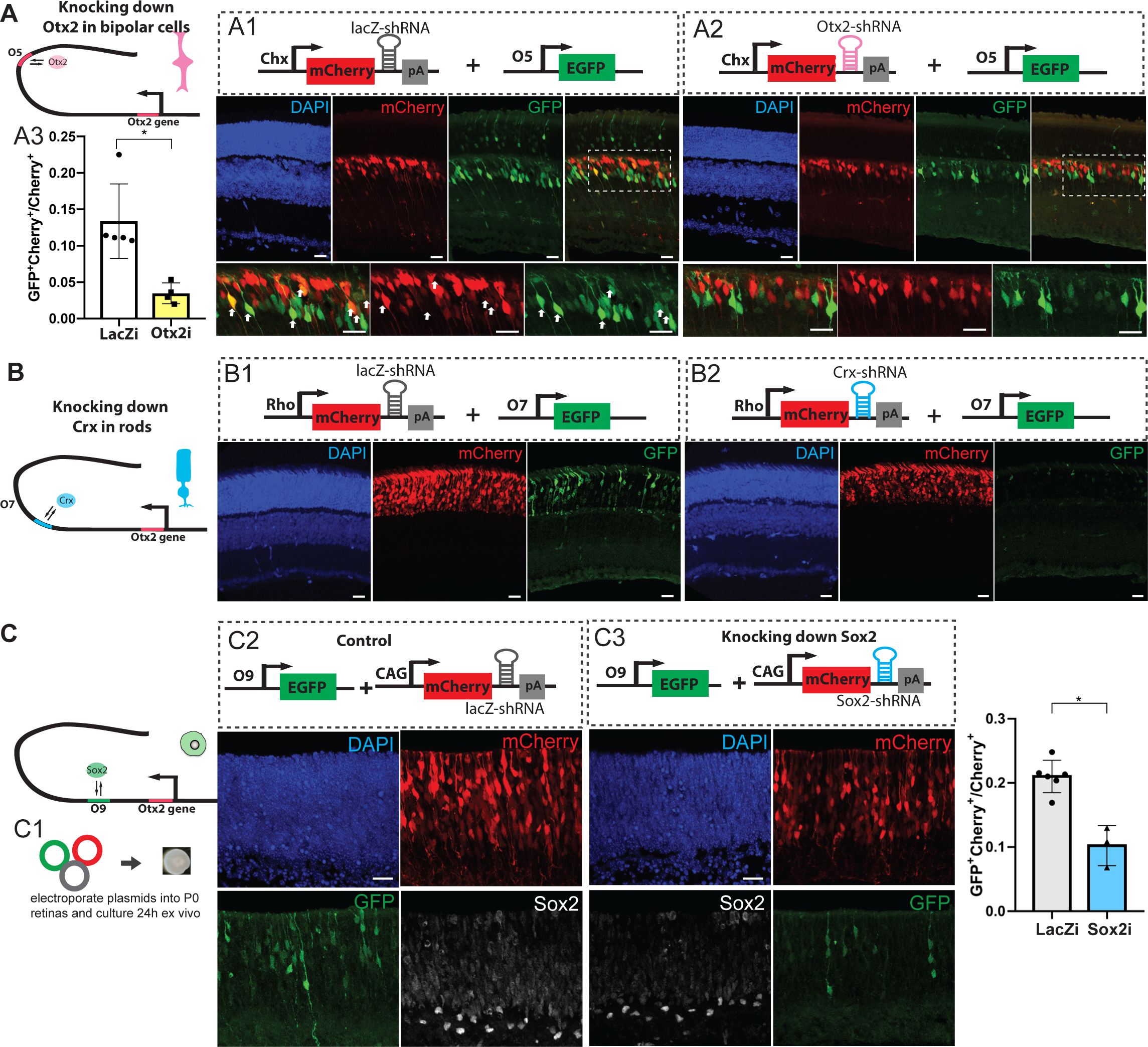
Otx2, Crx or Sox2 were required for the activation of O5, O7 or O9 CRM in the retina in vivo. A). Knocking down *Otx2* using shRNA diminished O5 activity in bipolar cells. Control LacZ-shRNA (A1) or Otx2-shRNA (A2) were placed in the 3’UTR of *mCherry*, and driven by the Chx164 CRM (Kim et al. 2008), which directs expression in mature bipolar cells. The shRNA-containing plasmid and O5-GFP plasmid were co-electroprated into retinas in vivo at P0. Retinas were harvested at P14. White arrows: mCherry+GFP+ positive cells. The percentage of mCherry+ cells that turned on the O5 CRM was quantified in A3. (B). Knocking down *Crx* in rods diminished O7 CRM activity. Control LacZ-shRNA (B1) or Crx-shRNA (B2) were placed in the 3’UTR of *mCherry*, and driven by the Rho promoter (Matsuda and Cepko 2007, 2004), which directs expression in mature rods. The shRNA-containing plasmid and O7-GFP plasmid were co-electroprated into retinas in vivo at P0. Retinas were harvested at P14. (C). Knocking down Sox2 diminished O9 activity in retinal cells at P1. Control LacZ-shRNA (C2-C3) or Sox2-shRNA (C4) were placed in the 3’UTR of *mCherry*, and driven by the ubiquitous CAG promoter. The shRNA-containing plasmids were electroprated into P0 retinas with the O9-GFP plasmid. The retinas were cultured ex vivo for 24 hours (C1). The percentage of mCherry+ cells that turned on the O9 CRM was quantified in C5. * P<0.01. T-test. Scale bar: 20um.

In mature retinas, Otx2 is expressed at higher levels in bipolar cells than in rods, which matches the pattern of O5-GFP expression (Figure 2C). Notably, compared with cone bipolar cells, rod bipolar cells express relatively lower levels of Otx2 (Supplementary Figure 6A-C). Consistent with this, the majority of O5-GFP+ bipolar cells had the morphology of cone bipolar cells (Figure 2, C2), suggesting that the level of Otx2 is important for the expression pattern of O5 CRM activity in bipolar cells. Interestingly, Crx is expressed at a higher level in rods than in bipolar cells (Glubrecht et al. 2009; Shekhar et al. 2016) (Supplementary figure 6B), and the O7-GFP, which requires a Crx binding site, was found to be active only in rods. It is possible that the level of Crx is not high enough to activate O7-GFP in bipolar cells. However, over-expression of Crx specifically in bipolar cells via the Chx164 bipolar CRM (Kim et al. 2008), did not lead to activation of O7-GFP in bipolar cells (data not shown). Similarly, Sox2 is expressed in Müller glial cells in the adult retina (Surzenko et al. 2013), but the O9-GFP CRM, which requires the Sox2 binding site, was not active in these cells at P14. We compared the levels of *Sox2* in P3 retinal cells with those in P14 Müller glial cells by quantitative smFISH (Supplementary figure 6F-G). The mRNA level of *Sox2* in mature Müller glial cells was found to be only slightly lower than that in RPCs at P3, suggesting that the level of *Sox2* is not responsible for the loss of O9 activity in mature Müller glial cells. It is possible that unknown transcriptional repressors of O7 or O9 CRMs exist in mature bipolar or Müller glial cells, a hypothesis that needs to be investigated.

**Figure 6.**
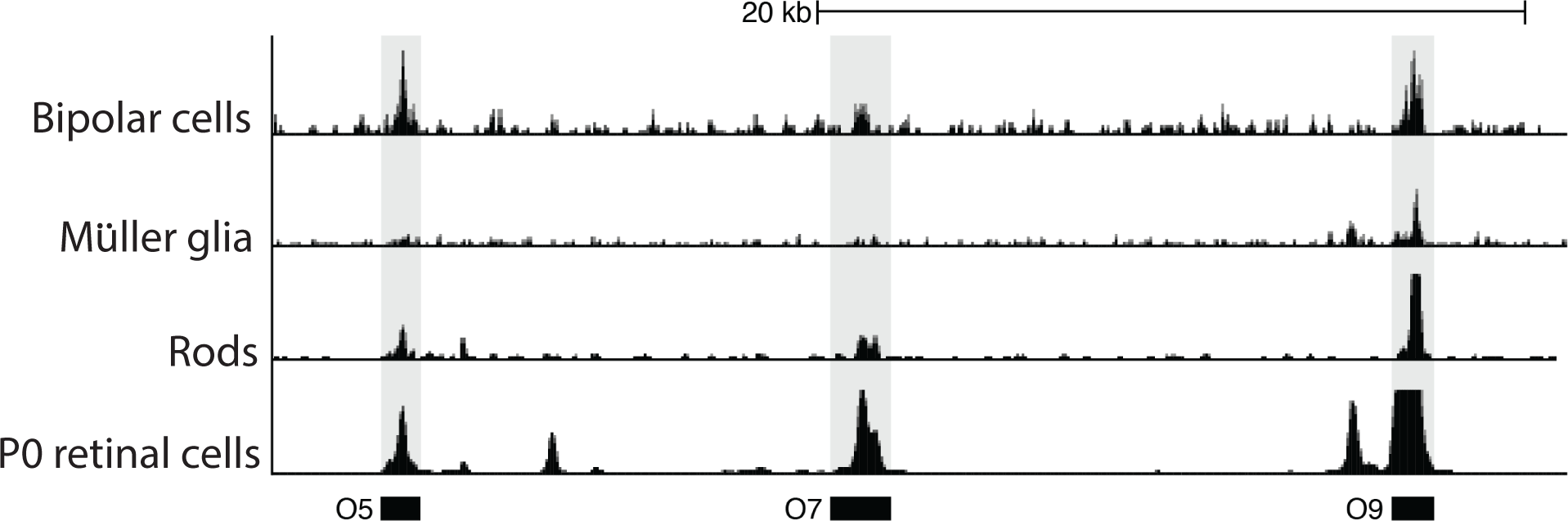
The chromatin status of O5, O7 and O9 CRMs in different retinal cell types and at different stages. The published ATAC-seq (bipolar, rod and Müller glial cells) (Hughes et al. 2017; Jorstad et al. 2017) and DNase-seq (P0 retinal cells) (Vierstra et al. 2014) results were re-analyzed to reveal the chromatin status of O5, O7 and O9 CRMs. All three CRMs were open and accessible in P0 retinal cells. The O5 CRM was very accessible in mature bipolar cells, poorly accessible in rods, and inaccessible in Müller glia. The O7 CRM was open in both bipolar cells and rods, but not in Müller glia. The O9 CRM was accessible in P0 retinal cells, bipolar cells and rods, and relatively less accessible in Müller glia.

Taken together, O5, O7, or O9 CRMs are active in cells expressing Otx2, Crx or Sox2, but not all Otx2, Crx or Sox2 positive cells turn on the corresponding CRMs. The levels of TFs appear to be important for CRM activity at neonatal stages. In the differentiated retina, the activity of repressors, particularly in mature bipolar and Müller glia with respect to the O7 and O9 CRMs, may contribute to the regulation of the CRM activity.

### Otx2, Crx and Sox2 are required for the activation of O5, O7 and O9 CRM in the retina

To investigate whether Otx2, Crx and Sox2 are required for O5, O7 and O9 activities in different retinal cell types at different stages in vivo, we used shRNA constructs to knock down these TFs. To test the necessity of Otx2 in bipolar cells for O5 activation, *Otx2* was knocked down using a miRNA-based shRNA cassette, which was inserted into the 3’ UTR of the *mCherry* gene (Wang et al. 2014). To specifically target differentiated bipolar cells, we used the bipolar specific CRM, 164bp Chx10 CRM (Chx164) (Kim et al. 2008). Even though the Chx164 CRM is biased towards labeling rod bipolar cells, i.e. less active in cone bipolar cells, it is the most effective driver for specific targeting of differentiated bipolar cells, and not RPCs, where Chx10 is also expressed. The Chx164-mCherry-Otx2-shRNA plasmid or control Chx164-mCherry-LacZ-shRNA plasmid was co-electroporated with the O5-GFP plasmid into P0 retinas in vivo. The retinas were harvested at P14 and the activity of the O5 CRM was assayed in cells marked by mCherry expression. The percentage of mCherry+GFP+ cells among mCherry+ cells was significantly reduced in the presence of *Otx2*-shRNA compared to the level in *LacZ*-shRNA controls (Figure 5, A3), suggesting that Otx2 is necessary for the activation of the O5 CRM in mature bipolar cells. Similarly, knocking down *Crx* in differentiated rods using the rod-specific rhodopsin promoter (Matsuda and Cepko 2007) significantly decreased the activity of the O7 CRM in vivo (Figure 5B). Knocking down *Sox2* in P0 retinas ex vivo greatly suppressed the activity of the O9 CRM (Figure 5C). These data demonstrate that Otx2, Crx and Sox2 are required for the activation of the O5, O7, and O9 CRMs, respectively.

### The chromatin status of *Otx2* CRMs in different retinal cell types

To further understand how multiple Otx2 CRMs collectively control *Otx2* transcription in the retina, we examined the chromatin status of endogenous O5, O7 or O9 CRMs in the genome in different retinal cell types and at different stages.

Heterochromatic regions of the genome, which are marked by H3K9me3 (Histone3 lysine9 trimethylation) or H3K27me3 (Histone3 lysine 27 trimethylation) (Becker et al. 2017), are typically resistant to DNaseI, while active or poised CRMs are hypersensitive to DNaseI. DNase-seq and ATAC-seq allow for genome-wide profiling of the “open” regions in the genome (Klemm et al. 2019). We reanalyzed the published DNase-seq and ATAC-seq data (Vierstra et al. 2014; Hughes et al. 2017; Jorstad et al. 2017), to ask whether the Otx2 CRMs were open or closed in retinal bipolar cells, rods, Müller glial cells, and P0 retinal cells (Figure 6). Genomic sequences corresponding to O5, O7 and O9 CRMs were found to be open in P0 retinal cells, which are composed of retinal progenitors/precursors and differentiated cells. In mature bipolar cells, the O7 CRM was less accessible, compared with the O5 and O9 CRMs. In rods, all three CRMs were found to be open. In Müller glial cells, the O5 and O7 CRMs were closed, and the O9 CRM was less accessible than in other cell types. Therefore, the endogenous O5, O7 and O9 CRMs have different chromatin accessibilities in different retinal cell types and at different stages during development.

## Discussion

We analyzed the regulation of *Otx2*, a gene required for the formation of several retinal cell types at different times in development. We wished to understand not only how its temporal and cell type specific transcription are orchestrated, but also how its level is controlled, given that its level can dictate whether a rod or a bipolar fate are chosen (Wang et al. 2014) as well as the survival and function of particular mature retinal cells (Bernard et al. 2014). We first used plasmid electroporation of genomic regions defined by their chromatin accessibility to identify several CRMs, O5, O7, and O9. These elements showed differential activity regarding cell types and the timing of their expression. The binding sites that are required for CRM activity were defined and were used to successfully direct a search for the cognate TFs. The identified TFs were found to be necessary, but not sufficient, for the expression patterns of these CRMs. This led us to investigate the accessibility of the CRMs in the chromatin of particular cell types in vivo. The O5, O7 and O9 CRMs were found to display differential chromatin accessibility in different cell types. Based on these data, we propose a model on how the stage- and cell type-specific expression of Otx2 is achieved in the retina (Figure 7A).

**Figure 7.**
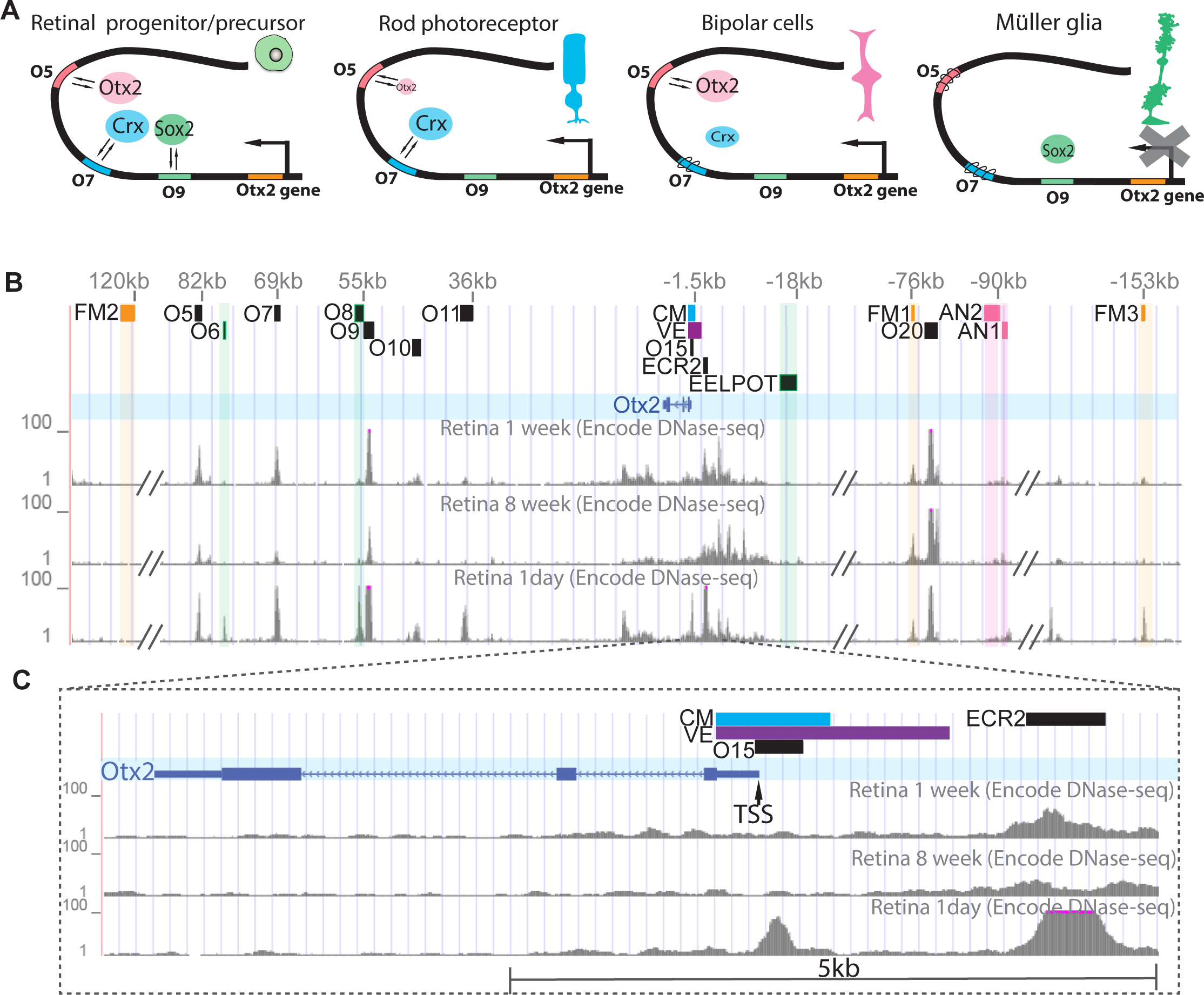
TFs and the chromatin status of CRMs collectively control the cell type- and stage-specific expression of Otx2 in the retina. **(A)** The proposed model. In retinal progenitor/precursor cells, O5, O7 and O9 CRMs are all active. The O9 CRM, which can be activated by Sox2, preferentially drives Otx2 expression in RPCs and their newly postmitotic progeny. Differentiated rods express high levels of Crx and low levels of Otx2. The O7 CRM, which can be activated by Crx, is the main CRM that is used to activate *Otx2* transcription in rods. Mature bipolar cells express relatively higher levels of Otx2 and lower levels of Crx. Apart from the reduced expression of Crx, the O7 CRM is less accessible in bipolar cells. The O5 CRM is thus the main CRM that drives Otx2 transcription in bipolar cells. While Müller glia express Sox2, which can activate the O9 CRM, Sox2 is not able to direct *Otx2* transcription in these cells, perhaps due to the presence of a repressor. Taken together, different transcription factors, their levels and the chromatin status of individual CRMs regulate the cell type- and stage-specific expression of the *Otx2* gene. **(B)** The CRMs regulating Otx2 expression in different tissues. The O5, O7, O9, O10, O11, O15, and O20 CRMs collectively regulated Otx2 expression in embryonic and postnatal retina. The EELPOT (Muranishi et al. 2011), O6 and O8 CRMs (Nadadur et al. 2019) were active in embryonic retina, but silenced in postnatal retina. AN1 and AN2: anterior neuroectoderm CRMs; FM1, FM2, and FM3: forebrain/midbrain CRMs; CM: cephalic mesenchyme CRM; VE: visceral endoderm CRM. **(C)** The high magnification view of the promoter region of the *Otx2* gene. TSS: transcription start site.

At the neonatal stage (P0-P3), the chromatin regions of O5, O7 and O9 CRMs are open. Their regulators, Otx2, Crx and Sox2 are present and collectively regulate the expression of Otx2 via these 3 CRMs in overlapping retinal cell populations. The levels of these TFs are important for CRM activity at this stage. In differentiated retinas (P14 or older), each CRM and its TF regulate *Otx2* expression in a distinct manner. In bipolar cells, the O5 and O9 CRMs, but not the O7 CRM, are significantly open. As Sox2 is not expressed in bipolar cells, the O9 CRM is not active in these cells. The expression of Otx2 in bipolar cells is self-regulated in a feed forward fashion via the O5 CRM. In rod photoreceptors, all three CRMs are open. The O9 CRM is not active in rods, likely due to the absence of Sox2. The O5 CRM is weakly active in rods likely due to the low expression of Otx2 in rods. Crx primarily regulates the transcription of *Otx2* via the O7 CRM in rod photoreceptors. In addition, Otx2 is not expressed by mature Müller glial cells. Even though Sox2 is present and the O9 CRM is open in these cells, the O9 CRM is not active in Müller glial cells, suggesting that Sox2 alone is not sufficient to activate the O9 CRM and Otx2 expression in Müller glial cells. Overall, retinal cells utilize the availability of TFs, levels of TFs, and the chromatin accessibility of multiple CRMs to precisely regulate Otx2 expression in a stage- and cell type-specific manner in the postnatal retina in vivo.

Otx2 also is expressed in the retina at embryonic stages (Muranishi et al. 2011). We examined the activities of the candidate Otx2 CRMs (Figure 1) in E14.5 (embryonic day 14.5) retinas ex vivo. Interestingly, all of the CRMs that are active in the postnatal retina, including O5, O7 and O9, were found to be active at E14.5. We recently used non-coding RNA (ncRNA) profiling to identify photoreceptor CRMs (Nadadur et al. 2019) and found that the O5, O7 and O9 CRMs overlapped with regions that showed transcription of ncRNA in the embryonic retina. Furthermore, regions of the chick genome with sequence homology to the mouse O5 and O7 CRMs showed CRM activity in the developing embryonic chick retina. Interestingly, as in mouse, Otx2 regulated the chick O5 CRM (Lonfat et al. in preparation). However, despite the activity of some CRMs at both embryonic and postnatal stages, a few *Otx2* CRMs active at embryonic stages were no longer active in the postnatal retina. The O6, O8 and EELPOT CRMs (Figure 7B)(Muranishi et al. 2011) are active in the embryonic retina, but are silent postnatally. Different mechanisms may control the developmental silencing of these CRMs. The O6 and O8 CRMs are DNaseI hypersensitive in the postnatal retina, and thus it is likely that it is the absence of their TFs, which are currently unknown, that leads to their lack of activity. The EELPOT CRM is not DNaseI hypersensitive in the postnatal retina (Figure 7B), and thus may not be accessible by its TFs, such as Rax, which is present at postnatal stages. These data suggest that additional CRMs contribute to the regulation of *Otx2* expression at embryonic stages, and that the silencing of embryonic CRMs at late stages is caused by chromatin remodeling and/or dynamic expression of TFs.

Apart from the retina, Otx2 is essential for early embryonic development and brain development (Beby and Lamonerie 2013; Acampora 1999). Several CRMs have been identified to control *Otx2* expression in visceral endoderm (VE), anterior neuroectoderm (AN), cephalic mesenchyme (CM) and developing forebrain/midbrain (FM) (Kurokawa et al. 2004a, 2004b, 2; Kimura et al. 2000; Bryan 1997). The published VE and CM CRMs are located in the promoter region of the *Otx2* gene, and partially overlap with the O15 CRM, which drives weak expression in the neonatal retina (Figure 1). Interestingly, none of the other distal *Otx2* CRMs are shared among tissues (Figure 7B). We examined the chromatin accessibility of the non-retinal *Otx2* CRMs in the retina to investigate whether this is due to differential chromatin states. The published anterior neuroectoderm CRMs (AN1 and AN2) (Kurokawa et al. 2004b) are not significantly open in the postnatal retina, while two out of three mouse forebrain/midbrain CRMs (FM1 and FM3, corresponding to O19 and O22) (Kurokawa et al. 2004a) are open, but inactive in the retina. This further suggests that the activities of the tissue-specific *Otx2* CRMs are not only regulated by differential chromatin states, but also due to the availability of their TFs.

Taken together, the expression of Otx2 is regulated via many distinct CRMs and TFs in different developmental and cellular contexts. Different tissues use non-overlapping distal CRMs to regulate Otx2 expression. Temporally, within the same tissue, such as the retina, some embryonically active CRMs are silent at postnatal stages, while all of the postnatally active CRMs can be active at embryonic stages. In addition to these temporal aspects, there are CRMs that preferentially regulate Otx2 expression in distinct cell types postnatally. Retinal progenitor/precursor cells, rod photoreceptors, and bipolar cells all express Otx2, but do so using a distinct combination of CRMs and TFs.

While improving our understanding of the complex regulation of *Otx2* gene, our study raises additional questions and future directions. First, how is the dynamic, differential accessibility of *Otx2* CRMs established in different tissues and cell types? The positioning and occupancy of nucleosomes are primary determinants of chromatin accessibility (Klemm et al. 2019). Poised and active CRMs reside in genomic regions with low nucleosome occupancy and high nucleosome turnover rate (He et al. 2010). TFs and chromatin remodelers are key regulators of the nucleosome occupancy and positioning. Given that chromatin remodelers lack sequence specificity, sequence-specific TFs are proposed to play central roles in establishing the differential accessibility of *Otx2* CRMs. It is not known whether the TFs that initiate remodeling of chromatin accessibility are the same TFs that activate CRMs. Otx2 and Sox2, which can activate the O5 and O9 CRMs in the retina, can function as pioneer factors and regulate chromatin remodeling in other systems (Iwafuchi-Doi and Zaret 2014; Boulay et al. 2017). Crx can regulate the chromatin remodeling of genes associated with photoreceptor differentiation (Ruzycki et al. 2018). It is highly possible that these tissue-or cell type-specific TFs help shape the chromatin accessibility, together with other components of the chromatin remodeling machinery. Consistent with this possibility, the O5 CRM is not open in Müller glial cells that do not express Otx2. Additional studies are needed to determine whether and how Otx2, Crx and Sox2 regulate the differential remodeling of chromatin accessibility in the retina.

Second, how are the differential expression levels of Otx2 established in different cellular contexts? Otx2 is expressed at a low level in rod photoreceptors relative to the high level in bipolar cells. Its expression level is essential for the binary cell fate decision of these two cell types, with high expression driving bipolar formation (Wang et al. 2014). One of the ways to establish the differential expression levels of Otx2 could be via chromatin accessibility and/or TF activity. For this decision, our data argue for the level of TFs as one of the key determinants, as when we increase the level of Otx2, Crx or Sox2, the activity of O5, O7 or O9 CRM in the neonatal retina is increased. These TF dose effects could help to establish and maintain the differential expression levels of Otx2 in rod and bipolar cells. Specifically, the O5 CRM is strongly active in bipolar cells, which express high levels of the O5 regulator, Otx2, creating a feedforward loop, which likely maintains high Otx2 levels in bipolar cells. In contrast, the level of Otx2 in rod photoreceptors is low, likely not high enough to stimulate the self-reinforcement mechanism. Instead, Crx, which is highly expressed in rods, regulates Otx2 expression via the O7 CRM. Additional factors, which titrate down *Otx2* levels in rod photoreceptor to prevent the feed-forward loop operating in bipolar cells, need to be identified to complete our understanding. Additional mechanisms, involving the Notch signaling pathway and TFs, including Blimp1 (Prdm1), Vsx2 and Sox2, contribute to the regulation of Otx2 expression in retinal progenitor cells and newly postmitotic cells (Muranishi et al. 2011; Wang et al. 2014; Kim et al. 2008). All of these factors will need to be integrated for the dynamic and specific regulation of Otx2 in different cellular contexts.

Third, what are the advantages of using multiple TFs and CRMs to regulate Otx2 expression in different cellular contexts? This strategy can significantly increase the robustness of the system. When one CRM element is mutated or disrupted, other CRMs could compensate for its function. For instance, when the O7 CRM was deleted by CRISPR/Cas9, we did not observe significant down regulation of *Otx2* transcription (data not shown). In addition, different cellular contexts probably maintain distinct regulatory environments. This strategy can ensure the activation of *Otx2* in response to different signals, to maximize the utility of TFs in the genome.

In addition to the data regarding Otx2 regulation provided by these studies, the CRMs identified here provide reagents for labeling and manipulating distinct cell types in the retina via in vivo plasmid DNA electroporation. The O5 CRM is primarily active in cone bipolar cells, the O9 CRM in neonatal retinal progenitor cells and newly postmitotic cells, and the O7 CRM marks rod photoreceptors.

In summary, we studied how the expression of a key regulator of retinal development, Otx2, is regulated, in order to understand the specifics of its expression. In addition, we wished to gain some insight into the mechanisms that might more generally account for the dynamic and cell type specific regulation of a gene. We show that multiple CRMs and a cohort of TFs collectively control the stage- and tissue/cell type-specific expression of Otx2. The chromatin accessibility of CRMs and the presence of sequence-specific upstream TFs are both required for activating *Otx2* expression in different cellular contexts. In the future, it will be important to investigate how the chromatin accessibility of CRMs are established and dynamically regulated across developmental and cellular states, and to understand how the TFs pools are maintained and regulated in different cellular contexts.

## Methods and Materials

### Animals

Wild type mouse neonates were obtained from time pregnant CD1 mice (Charles River Laboratories, #022). All animal studies were approved by Administrative Panel on Laboratory Animal Care (APLAC) at Stanford University.

### Plasmid Construction

The CAG-mCherry plasmid was from Matsuda (Matsuda and Cepko 2004). The Stagia3 vector was from Billings (Billings et al. 2010). The CAG-BFP plasmid was from Tang (Tang et al. 2015). All other plasmids were constructed by restrictive enzyme-based cloning or Gibson assembly methods (Gibson et al. 2009).

#### Otx2 CRM reporter plasmids (Figure 1)

The 25 candidate Otx2 CRMs were amplified from the mouse genome by primers showed in Table 1 and cloned into the Stagia3 vector. The sequences of these CRMs were shown in supplementary file 1.

#### CRISPR plasmids (Figure 3)

To generate the O5-Cas9 plasmid, the Cas9 fragment was amplified by PCR from Px330 plasmid (Addgene # 42230)(Cong et al. 2013) and replaced the GFP fragment in the O5-GFP plasmid. To generate the O5-sgRNA1 and O5-sgRNA2 plasmids, the following corresponding complementary oligonucleotides were annealed and cloned into the modified Px330 plasmid, which does not express Cas9.

O5-sgRNA1-f: 5’-TTG aacagccgacaaaacccgggGTTTAAGAGC

O5-sgRNA1-r: 5’-TTAGCTCTTAAACcccgggttttgtcggctgttCAACAAG

O5-sgRNA2-f: 5’-TTG gttctgcgggaccgccgttt GTTTAAGAGC

O5-sgRNA2-r: 5’-TTAGCTCTTAAACaaacggcggtcccgcagaacCAACAAG

#### CRM deletion analysis (Figure 4)

The truncated CRM O5-1 to O5-x were amplified by PCR and cloned into the Stagia3 vector plasmid individually. Mutation variants of O5-8, O7-1, or O9-12 were generated by annealing complementary oligonucleotides of individual mutated fragments and ligating them into the EcoRI and XhoI digested Stagia3 plasmid. The sequences of the truncated and mutated CRMs are shown in Supplementary file 1.

#### Cell type-specific knockdown and over-expression of Otx2, Crx or Sox2 (Figure 5)

The Chx-mCherry-lacZ-shRNA or Chx-mCherry-Otx2-shRNA plasmid was generated by replacing the CAG promoter of the CAG-mCherry-lacZ-shRNA (Addgene #73978, (Wang et al. 2014)) or CAG-mCherry-Otx2-shRNA (Addgene #73980, Wang et al. 2014) plasmid with the 164bp Chx10 CRM and basic TATA box promoter (Kim et al. 2008). The Rho-mCherry-lacZ-shRNA or Rho-mCherry-Crx-shRNA plasmid was generated by replacing the CAG promoter of CAG-mCherry-lacZ-shRNA (Addgene #73978, Wang et al. 2014) or CAG-mCherry-Crx-shRNA (Addgene #73987, (Wang et al. 2014) plasmid with the 2.2kb bovine rhodopsin promoter (Matsuda and Cepko 2007). The RNAi cassette against Sox2 gene was obtained via the BLOCK-iT™ PolII miR RNAi Express (Life Technologies) and inserted into the RNAi expression vector CAG-mCherry-miR155 (Gift from Dr. Jianming Jiang, National University of Singapore). The sequence of the Sox2 RNAi cassette is: 5’TGCTGTGGTCATGGAGTTGTACTGCAGTTTTGGCCACTGACTGACTGCAGTACCT CCATGACCA.

The CAG-Sox2 plasmid was generated by cloning the ORF of mouse *Sox2* gene into the CAG-GFP backbone (Addgene #11150, (Matsuda and Cepko 2004). The GFP fragment was replaced by the *Sox2* ORF. The CAG-filler-DNA plasmid was generated by deleting the GFP fragment from the CAG-GFP plasmid.

### In vivo and ex vivo plasmid electroporation into the retina

Ex vivo and in vivo retina electroporations were carried out as previously described (Matsuda and Cepko 2004, 2007; Wang et al. 2014). For ex vivo electroporation, 5 pulses of 25V, 50ms each and 950ms interval were applied to dissected retinas. For in vivo electroporation, 5 pulses of 80V, 50ms each and 950ms interval were applied to neonatal mouse pups. All ex vivo and in vivo electroporation experiments were repeated with at least three biological replicates. Plasmids were electroporated with a concentration of 500ng/ul to 1ug/ul per plasmid.

### Histology and immunohistochemistry

Dissected mouse eyeballs were fixed in 4% PFA in 1XPBS (pH7.4) for 2 hours at room temperature. Retinas then were dissected and equilibrated at room temperature in a series of sucrose solutions (5% sucrose in 1XPBS, 5min; 15% sucrose in 1XPBS, 15min; 30% sucrose in 1XPBS, 1 hour; 1:1 mixed solution of OCT and 30% sucrose in PBS, 4 degree, overnight), frozen and stored at −80°C. A Leica CM3050S cryostat (Leica Microsystems) was used to prepared 20um cryosections. Retinal cryo-sections were washed in 1XPBS briefly, incubated in 0.2% Triton, 1XPBS for 20 min, and blocked for 30 min in blocking solution, which is 0.1% Triton, 1% BSA and 10% Donkey Serum (Jackson ImmunoResearch Laboratories, Inc.) in 1XPBS. Slides were incubated with primary antibodies diluted in blocking solution in a humidified chamber at room temperature for 2 hours or at 4°C overnight. After washing in 0.1% Triton 1XPBS three times, slides were incubated with secondary antibodies and DAPI (Sigma-Aldrich; D9542) for 30min to 2 hours, washed three times with 0.1% Triton, 1XPBS and mounted in Fluoromount-G (Southern Biotechnology Associates).

The following primary antibodies were used: chicken anti-GFP (Abcam, AB13970, 1:1000), goat anti-RFP (Fisher Sicentific, NC1578084, 1:500), rabbit anti-Otx2 (VWR, 10091-640, 1:1000), rabbit anti-Crx (Fisher Scientific, NBP215964, 1:500), mouse anti-Ki67 (BD Biosciences, 550609, 1:200), sheep anti-Chx10 (Exalpha Biologicals, X1179P, 1:500), and mouse anti-Sox2 (Santa Cruz Biotechnology, sc-365823) antibodies. EdU detection was performed with a Click-iT EdU Alexa Fluor 647 imaging kit (C10340, Invitrogen). AP activity was detected by an AP detection kit (Sigma, SCR004).

### Dissociation of retina tissues and quantitative single molecule FISH

Retinal tissues were dissociated as described previously (Trimarchi et al. 2008). After dissociation, retinal cells were left on PDL (Poly-D-Lysine, 0.1mg/ml, Millipore) treated slides for 45 min at 37°C. Retinal cells were then fixed on the slides in 4% PFA in 1XPBS (pH7.4) for 15min at room temperature. The cells were stained immediately or dehydrated by going through the 50%, 70% and 100% ethanol gradient, and stored in 100% ethanol at −20°C. FISH probes were purchased from RNAscope. The commercial protocol provided by RNAscope was followed. The slides were imaged by Zeiss confocal LSM 880. Images were analyzed by Zen software (Zeiss), and quantified by Fiji.

### EMSA assay

EMSA assays were performed as described previously (Wang et al. 2014). Roughly 1X10^6^ 293T cells were transfected with 1 μg of CAG-LacZ, CAG-Otx2, CAG-Crx, or CAG-Sox2 plasmid using PEI (1mg/ml, 4ul per 1μg DNA, 9002-98-6, Polysciences Inc). Nuclear extracts were prepared from these cells or P3 mouse retinas using NE-PER nuclear and cytoplasmic extraction reagent kit (Pierce). Complementary oligonucleotides were ordered, annealed and labeled by DNA 3’ End Biotinylation kit (Pierce). Chemiluminescent Nucleic Acid Detection Module (Pierce) was used to detect biotin-labeled probes after EMSA assay.

### Imaging and analysis

All images of retinal sections were acquired by a Zeiss LSM880 inverted confocal microscope. Retina explants were imaged by Leica M165 FC microscope. HEK293T cells were imaged by Zeiss Axio Observer 3 microscope. Images in Figure 2, 3 and 5 were maximum projections of 5 μm tissues and were quantified by Fiji software.

### TF binding sites prediction

To predict TF binding sites (TFBSs) located in CRMs, the Position Weight Matrix (PWM) annotated by MotifDB was used (Shannon, P. MotifDb: An Annotated Collection of Protein-DNA Binding Sequence Motifs v1.12.1, The R Foundation, 2015). Based on these PWM, FIMO (Grant et al. 2011) were used to search for potential TFBSs with the default cutoff E-value < 10^−4^.

### DNase-seq and ATAC-seq and data analysis

We downloaded the BigWig file for DNase-seq of mouse newborn retina tissue from the ENCODE portal (https://doi.org/10.1093/nar/gkx1081) (https://www.encodeproject.org/) with the following identifier: ENCSR000CNV (Vierstra et al. 2014). BigWig for rod photoreceptors was from (Hughes et al., 2017) and downloaded from GEO (Sample GSM2199333). Raw ATAC-seq data for bipolar cells and Mueller glia were from Jorstad et al. 2017 and downloaded from NCBI Sequence Read Archive (accession SRX2881305 and SRX2881310 respectively). Briefly, reads were aligned to mouse mm10 genome using Bowtie2 (-X 2000) (Langmead and Salzberg, 2012) and processed with SAMtools to remove duplicates and MT reads (Li et al., 2009). We generated BigWig coverage tracks using deepTools bamCoverage (Ramírez et al. 2016 doi:10.1093/nar/gkw257) with the following parameters: --binSize 1 -p=max --normalizeUsing RPGC -- effectiveGenomeSize 2308125349. BigWig for all datasets were visualized in UCSC Genome browser.

## Acknowledgements

Support was provided by American Diabetes Association and P30 to Stanford Ophthalmology (to S.W.) and Howard Hughes Medical Institute (to C.L.C). N.L. was supported by post-doctoral fellowships from the Swiss National Science Foundation and the Human Frontiers Science Program and by R01EY029771. The members of the Cepko and Wang laboratories provided valuable discussion and support for this project.

## Author contributions

S. W. and C. Chan designed the study, performed and analyzed most of the experiments, and wrote the manuscript. N. L. analyzed the DNase-seq and ATAC-seq data, and wrote the manuscript. S. Z. generated maxi preps for some of the plasmids used in this manuscript. A. D., L. L., M. W. and C. L. collected mouse experimental data for CRM screen. Z. J. performed the TFBS bioinformatic analyses. C. Cepko and S. W. designed and supervised the study, and wrote the manuscript. All authors discussed the data and commented on the manuscript.

